# Discovering cryptic splice mutations in cancers via a deep neural network framework

**DOI:** 10.1101/2022.10.14.512264

**Authors:** Raphaël Teboul, Michalina Grabias, Jessica Zucman-Rossi, Eric Letouzé

## Abstract

Somatic mutations can disrupt splicing regulatory elements and have dramatic effects on cancer genes, yet the functional consequences of mutations located in extended splice regions is difficult to predict. Here, we use a deep neural network (SpliceAI) to characterize the landscape of splice-altering mutations in cancer. In our in-house liver cancer series, SpliceAI uncovers many cryptic splice mutations, located outside essential splice sites, that validate at a high rate in matched RNA-seq data. We then extend the analysis to a large pan-cancer cohort of 18,115 tumors, revealing >100,000 cryptic splice mutations. Taking into account these mutations increases the power of driver gene discovery, revealing >100 new candidate driver genes. It also reveals new driver mutations in known cancer genes, doubling the frequency of splice alterations in tumor suppressor genes. Mutational signature analysis reveals the mutational processes that give rise to splice mutations in each cancer type, with an enrichment of signatures related to clock-like processes and DNA repair deficiency. Altogether, this work sheds light on the causes and impact of cryptic splice mutations in cancer, and highlights the power of deep learning approaches to better annotate the functional consequences of mutations in oncology.

## INTRODUCTION

RNA splicing is a key process in biology where non-coding regions of RNA (introns) are removed and exons joined together, to generate the mature messenger RNA (mRNA) that will be translated (1). In eukaryotes, alternative splicing occurs in ~95% of multi-exonic genes and generates a diversity of protein isoforms from a single gene (2). This process provides evolutionary flexibility and may have contributed to the development of multicellular organisms (3). In humans, splicing involves a series of reactions catalyzed by a large ribonucleoprotein complex called the spliceosome, which recognizes key regulatory sequences on the pre-mRNA. The two bases at each extremity of the intron constitute the essential donor (GU) and acceptor (AG) sites. Other cis-regulatory elements include the polypyrimidine tract and the branch point located around 30 nucleotides upstream the acceptor site, as well as exonic and intronic splicing enhancers and silencers (4). Yet, these sequences are less evolutionary conserved than the essential splice sites. RNA splicing is thus carefully regulated, and alterations of the splice machinery components or cis-regulatory sequences underlie many diseases (5–7). In cancer, core spliceosome components including *SF3B1, SRSF2* or *U2AF1* are recurrently altered in both hematological and solid tumors (8, 9). In addition, splice mutations are a frequent mechanism leading to the inactivation of tumor suppressor genes, and have been reported in 13% of genes in the Cancer Gene Census (9, 10). Mutations affecting essential splice sites lead to various degrees of intron retention or exon skipping and are thus annotated as damaging in cancer genomics projects. In contrast, the functional consequences of mutations in extended splice regions is difficult to predict because other regulatory elements largely involve degenerate sequence features. Several computational methods have been developed to automatically detect splice sites in DNA sequences and to predict the splicing impact of mutations. One of the earliest tools for detecting splice sites in eukaryotic mRNA was a decision tree via maximal dependence decomposition (11). This idea was enhanced in GeneSplicer by Markov models that capture additional dependencies among neighboring bases in splice regions (12). Other methods were introduced to model splice sites with Bayes networks (13) or pairwise correlations (14). Yeo and Burge proposed the MaxEntScan framework modeling sequence motifs based on the maximum entropy principle (15). This tool was shown to outperform previous probabilistic models and is used by variant annotation tools like the Variant Effect Predictor (VEP) (16). Neural networks were introduced to identify splice sites in NNSplice (17). More recently, a 32-layer deep residual neural network (SpliceAI) was developed to predict whether each position in a pre-mRNA transcript is a splice donor, acceptor or neither (18). In contrast to previous methods that only consider short windows around exon-intron boundaries, SpliceAI uses 10,000 nucleotide sequences as input, allowing to capture long-range determinants of splicing. This tool largely outperformed previous methods and was successfully used to unravel pathogenic cryptic splice mutations in patients with rare genetic diseases.

Here, we use a large series of liver tumors with matched whole genome/exome sequencing and RNA sequencing data to benchmark SpliceAI in a cancer setting. We then apply this tool to uncover cryptic splice mutations in a large pan-cancer series totaling >17,000 tumors. We show that SpliceAI outperforms previous widely used tools, and reveals numerous cryptic splice mutations that enhance the landscape of cancer driver genes and mutations.

## MATERIAL AND METHODS

### Datasets

We used an in-house series of 401 liver cancers with matched whole exome (WES, n=275) / genome (WGS, n=126) sequencing and RNA-seq data to benchmark the performance of SpliceAI. Initial bioinformatic analyses (alignment and variant calling) were already performed in previous studies (19–25) and we directly used processed data in this study. We then analyzed a large pan-cancer series comprising 17,714 tumor samples across 25 cancer types from the ICGC-PCAWG (n=2,699 WGS) (26), Hartwig Medical Foundation (n=4,818 WGS) (27) and TCGA (n=10,197 WES) (28) projects. Somatic mutations were downloaded directly from the online repositories of each project.

### Predicting splice-affecting mutations with SpliceAI

*We* used SpliceAI (v1.3.1) to predict the splicing impact of almost 300 millions somatic mutations. This computation-intensive task was performed on the Jean Zay supercomputer, the world’s 10th most powerful supercomputer, with a computing power reaching 28 petaflops. Computations were heavily performed on a GPU V100 allocation which allowed to shrink the computation time down to 0.003 seconds. SpliceAI returns 4 delta scores (DS) for each type of splicing abnormality (donor loss/gain, acceptor loss/gain). We retrieved for each mutation the strongest DS, the corresponding splicing consequence and its position relative to the mutation. Mutations with a DS ≥ 0.5 were considered to have a significant impact on splicing, as suggested in the original publication (18).

### Validation of splice-affecting mutations using matched RNA-seq data

Let *p* be the position of a predicted abnormal splicing event. Let *R_c_* be the set of reads crossing position *p* in a control sample with a MapQ score ≥ 20 and *R_m_* be the set of reads crossing position p in the mutated sample of interest with a MapQ score ≥ 20. We define *M_n_* and *M_α_* as a partition of *R_m_* such as *R_m_* = *M_n_* ∪ *M_α_*. We define *C_n_* and *C_α_* as a partition of *R_c_* such as *R_c_* = *C_n_* ∪ *C_α_*. *M_α_*(respectively *M_n_*) represents the ensemble of abnormal (respectively normal) reads in the mutated sample of interest. *C_α_* (respectively *C_n_*) represents the ensemble of abnormal (respectively normal) reads in the control samples. We calculated the Relative Usage of the Novel Junction (RUNJ) as previously described (18):

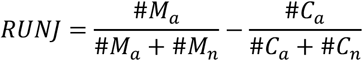

Two custom control panels were used for each mutation: a control PON (Panel of Normal) composed of 5 non-tumor samples from the same sequencing series as the sample of interest, and a control POT (Panel of Tumors) composed of 5 tumor samples from the same sequencing series and without a somatic mutation at the same locus. RUNJ scores were calculated in two ways, considering as abnormal reads only those supporting the most direct consequence of the splicing alteration (e.g. reads passing trough the essential donor for donor loss events) in the RUNJ_direct_ score, or all reads supporting abnormal junctions in the RUNJ_direct_ score. Finally, a prediction was validated if and only if all of the following hypotheses are verified:

- *#M_α_* ≥ 2
- *RUNJ_direct,PON_* ≥ *lim* or *RUNJ_direct,PON_ ≥ lim*
- *RUNJ_all,POT_* ≥ *lim* or *RUNJ_all,POT_* ≥ *lim* or not applicable.

To facilitate the validation of large numbers of variants, we created the *PyRNA* library that allows retrieving relevant reads from RNA-seq bam files and calculating RUNJ scores: https://github.com/FunGeST/pyRNA.

### Benchmarking the performance of SpliceAI against GeneSplicer and MaxEntScan

We used the two widely used tools GeneSplicer (12) and MaxEntScan (15) to benchmark the performance of SpliceAI in predicting splice-affecting mutations. We downloaded the MaxEntScan and GeneSplicer softwares from http://genes.mit.edu/burgelab/maxent/download/ and http://www.cs.jhu.edu/genomics/GeneSplicer/ respectively. Both tools were run with default parameters to predict the impact on splicing of the 1,704,461 somatic mutations in th LICA-FR series. For comparison, we applied thresholds for GeneSplicer and MaxEntScan scores in order to have the same number of positive predictions as SpliceAi for three DS thresholds (0.2, 0.5, 0.8). We validated the predictions of the 3 tools using matched RNA-seq data as described above. We computed the proportion validated predictions and the area under the precision-recall curve (PR-AUC) for each tool.

### Driver gene analysis

We used MutSigCV to identify genes with significantly more mutations than expected by chance considering mutation categories and genomic covariates, as previously described (29). For each tumor series, we first run MutSigCV with classical categories of coding mutations (missense, nonsense, essential splice, frameshift, inframe indels or synonymous). We then re-run the analysis after converting to splice variants the cryptic splice mutations uncovered by SpliceAI (DS ≥ 0.5). We then compared MutSigCV q-values obtained with both settings to identify new genes reaching significance when adding cryptic splice variants. We also calculated the mutation frequency of each gene in the Cancer Gene Census (10) considering only classical categories of coding mutations or adding cryptic splice sites.

### Mutational signature analysis

We used Palimpsest (30) to quantify the contribution of known COSMIC signatures in 2,699 tumors of the ICGC PCAWG data set. To ensure that only relevant signatures were included, we analyzed each cancer type separately and we included only signatures that were shown to be present in that cancer type (31). We then used Palimpsest to quantify the probability *p_m,sig_* of each mutation *m* being due to each mutational process with signature *sig*, as previously described (20). Finally, we compared the contribution of signatures to splice mutations and other mutations in each series. The contribution of signature *sig* to a set of mutations *S_m_* was estimated as the sum:

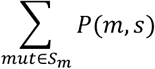

To identify mutational processes preferentially giving rise to splice mutations, we compared the distribution of probabilities *P(m,s)* for each signatures *s* between splice mutations and others using Wilcoxon rank sum tests.

## RESULTS

### SpliceAI outperforms traditional tools for predicting cryptic splice mutations in cancer

We used an in-house series of 401 liver cancers (19–25) with whole exome (n=275) or whole genome sequencing (n=126) and matched RNA-seq data to predict and validate cryptic splice mutations using SpliceAI, MaxEntScan and GeneSplicer (**Figure 1A**). A total of 1,704,461 somatic mutations were analyzed with the three tools. SpliceAI returned predictions for 432,773 mutations, covering most intragenic mutations. By contrast, GeneSplicer returned predictions for only 13,589 mutations, and MaxEntScan 6,317 mutations, mostly located in splice regions. SpliceAI provides for each mutation a delta score (DS) representing the probability that a mutation creates (positive DS) or disrupts (negative DS) a splice site. 2,056 mutations were predicted to alter splicing with an absolute DS ≥ 0.5 (threshold recommended by SpliceAI developers), including 649 acceptor loss (AL), 753 donor loss (DL), 263 acceptor gain (AG) and 391 donor gain (DG) events (**Supplementary Table S1** and **Supplementary Figure S1**). Mutations with an absolute DS ≥ 0.9 were strongly enriched in essential splice sites (67%). By contrast, 60% of mutations with an absolute DS ≥ 0.5 were located outside canonical splice sites (**Figure 1B**). For comparison, we applied thresholds on GeneSplicer and MaxEntScan scores so that the number of mutations predicted to impact splicing was equivalent for the 3 tools. We then used matched RNA-seq data to validate the predictions. For each mutation predicted to affect splicing, we identified reads consistent with abnormal vs. normal splicing. We then calculated the ratio of abnormal reads over the total number of reads (UNJ score), which we normalized against UNJ scores in a series of normal and non-mutated tumor samples to obtain the RUNJ scores (**Figure 1C**). Mutations with a RUNJ ≥ 0.01 and at least 2 reads supporting the abnormal splicing events were considered validated. 357 mutations were predicted to alter splicing by all the methods and showed the best validation rate (90%, **Figure 1D**). SpliceAI and MaxEntScan had more consistent predictions, with 1,159 mutations predicted to impact splicing by both tools but not GeneSplicer. Among tool-specific mutations, SpliceAI-only mutations showed the best validation rate (64 % vs. 41% for MaxEntScan & 31% for GeneSplicer). The precision-recall curve shows that the best overall performance was achieved with SpliceAI, with a PR-AUC=0.54 (**Figure 1E**). Two examples of cryptic splice mutations affecting liver cancer driver genes uncovered by SpliceAI are shown in **Figure 1**. These include a TP53 mutation initially annotated as synonymous that actually disrupts the adjacent splice donor site, leading to various degrees of intron retention (**Figure 1F**). Another interesting example is a missense mutation in the tumor suppressor gene ARID1A that creates an acceptor gain 3 bases downstream. RNA-seq reads show that this cryptic acceptor is used in 12% of the transcripts, leading to a dual consequence of the mutation: missense or frameshift depending on the transcripts (**Figure 1G**). Altogether, these data indicate that SpliceAI outperforms previous tools for the detection of splice-impacting mutations, and can reveal new cryptic splice mutations in cancer driver genes.

**Figure 1.**
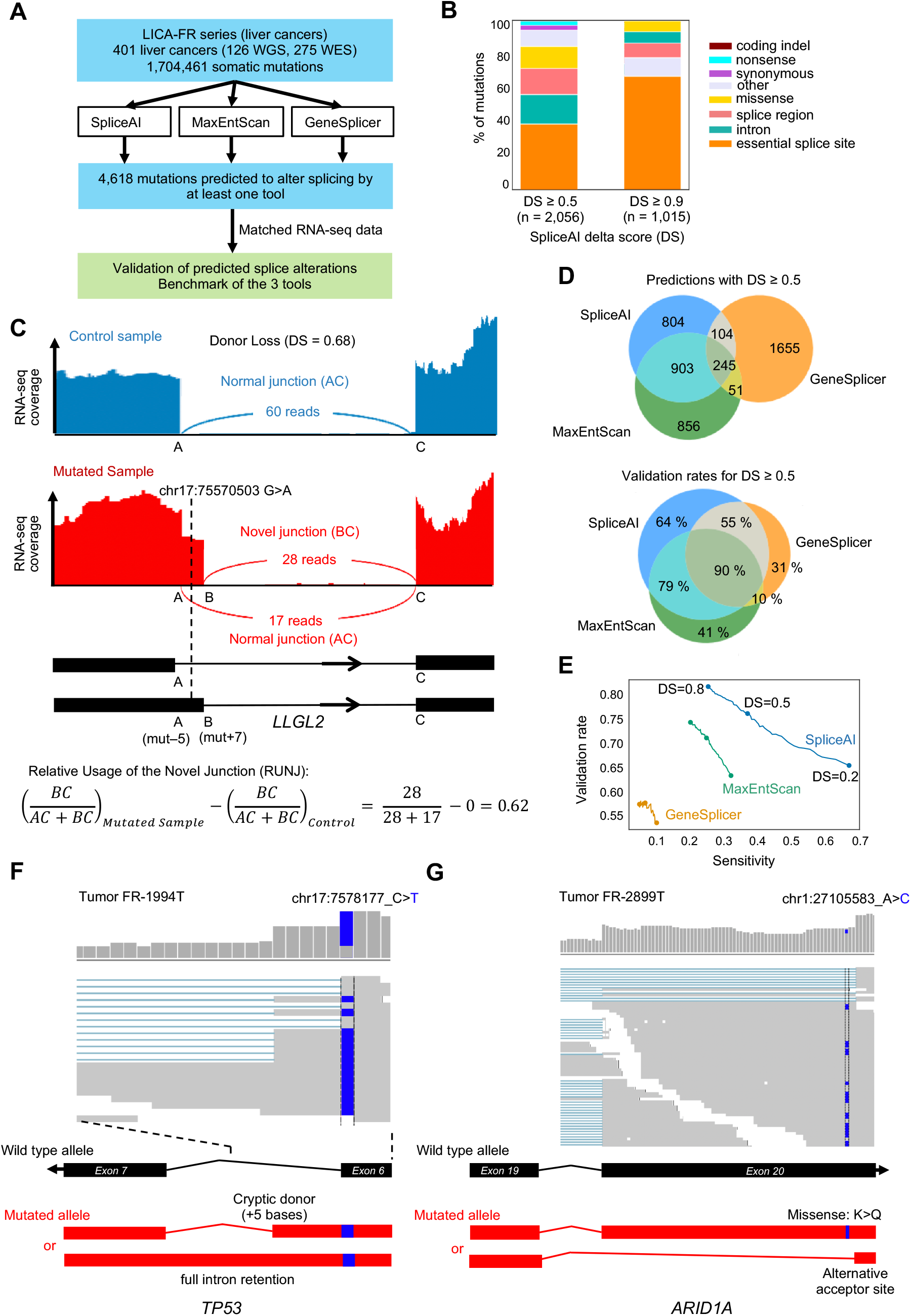
Benchmark of splice prediction tools on the LICA-FR dataset. (**A**) Schematic diagram describing the benchmark study. (**B**) Proportion of initial mutation categories for mutations predicted to impact splicing by SpliceAI with different delta scores (DS). (**C**) Example of RNA-seq validation of a predicted cryptic splice mutation. (**D**) Venn diagrams showing the number of overlap of mutations predicted to alter splicing by the 3 tools (top) and the proportion of mutations validated in matched RNA-seq data (bottom). (**E**) Validation rate and sensitivity obtained with the 3 tools for different thresholds on prediction scores. (**F**) Example of cryptic splice variant in TP53 initially annotated as synonymous mutation. (**G**) Example of cryptic splice variant in ARID1A initially annotated as missense mutation.

### Pan-cancer analysis of cryptic splice mutations

We next applied SpliceAI to uncover cryptic splice mutations in a large compendium of 17,714 tumor samples across 25 cancer types from the ICGC-PCAWG (n=2,699 WGS, mostly primary tumors (26)), Hartwig Medical Foundation (n=4,818 WGS of metastatic cancers (27)) and TCGA (n=10,197 WES, mostly primary tumors (28)) projects (**Figure 2A**). To analyze the 277,415,611 somatic mutations, we used the Jean Zay supercomputer V100 quadri GPU partition, which took 0.003 second per variant to compute a prediction. We identified 203,889 mutations predicted to impact splicing with a DS ≥ 0.5, with a predominance of donor/acceptor losses (71.5%) over donor/acceptor gains (28.5%, **Figure 2B**). Of these, 43.5% corresponded to essential splice sites whereas 56.5% were cryptic splice mutations located in extended splice regions (13%), introns (14%) or exons (26%, **Figure 2C,D**). In addition to the two first intronic bases corresponding to essential splice sites, donor loss mutations frequently involved the last exonic base (21%) and the 5th intronic base (9%), indicating an important role of these positions in regulating the recognition of donor sites (**Figure 2E**). Interestingly, donor gain mutations also frequently involved the same positions (0 and +5, 9%, **Figure 2F**). Thus, by altering these bases that are important for the regulation of the native donor sites, these mutations favor the usage of alternative cryptic donors. Cryptic acceptor loss mutations were mostly located between the 3rd and 15th intronic bases (17%), corresponding to the polypyrimidine tract. Acceptor gains were quite evenly distributed between positions −15 and +3 relative to the closest essential acceptor site.

**Figure 2.**
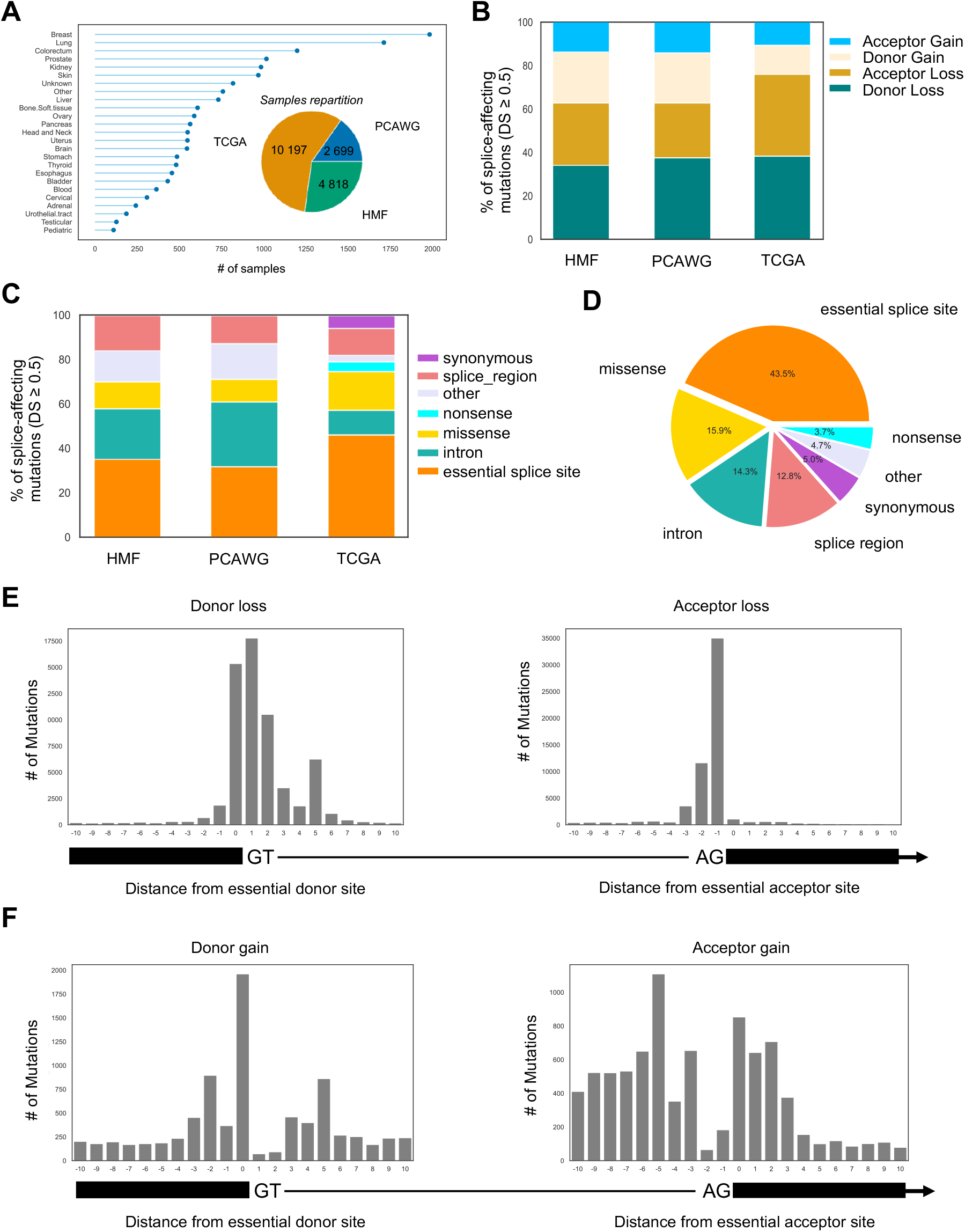
Pan-cancer analysis of cryptic splice mutations with SpliceAI. (**A**) Number, origin and cancer type of samples analyzed. (**B**) Proportion of splice altering mutations identified in each series classified by type of alteration (DL: donor loss, AL: acceptor loss, DG: donor gain, AG: acceptor gain). (**C**) Proportion of initial mutation categories for mutations predicted to impact splicing by SpliceAI in each cohort (DS > 0.5). (**D**) Overall proportion of initial mutation categories for the 203,889 mutations predicted to impact splicing with a DS > 0.5 in the whole data set. (**E**) Position of donor and acceptor loss mutations identified by SpliceAI relative to the closest essential splice acceptor and donor. (**F**) Position of donor and acceptor gain mutations identified by SpliceAI relative to the closest essential splice acceptor and donor.

### Cryptic splice mutations enhance the landscape of driver genes and mutations

We next investigated whether considering cryptic splice mutations would uncover new cancer driver genes and new damaging mutations in known cancer genes. To address this point, we used MutSigCV (29) to identify genes with significantly recurrent mutations in the HMF and TCGA series, considering classical VEP annotations or adding cryptic splice mutations. MutSigCV identified a median of 13.5 significantly mutated genes per series (range 1-121) with classical annotations. Including cryptic splice mutations added a total of 136 significant genes (median=3.5 per series, range=1-16, **Figure 3A**, **Supplementary Table S2** and **Supplementary Figure S2**). Extra significant genes were enriched in known cancer genes (n=38, 28%, P=3.0e-24), validating the biological relevance of the new findings. For example, *ARID1A*, a known driver gene in cholangiocarcinoma, was not significant in the relatively small TCGA_CHOL series (n=36) but became significant when adding cryptic splice mutations. This was also the case for *SMAD4* in pancreatic cancer, *TBX3* in colorectal cancer, or *TSC2* in liver cancer. Adding cryptic splice mutations also revealed extra-significant genes known to be drivers in other cancer types. For example, we identified in the HMF breast metastasis series significantly recurrent mutations of *CDKN1A* (known driver in bladder cancer), *MAX* (known in paragangliomas, endometrioid and colon carcinomas) and *CDK12* (known in ovarian cancer). Finally, 98 of the genes that became significant when adding cryptic splice mutations were not in the Cancer Gene Census and may correspond to new driver genes. These included *GPS2*, negative regulator of RAS- and MAPK-mediated signal (32), recurrently mutated in prostate cancer; *BRD7*, member of the SWI/SNF chromatin remodeling complex, recurrently altered in urothelial cancer; *GPAM*, associated with steatosis and liver damage (33), recurrently mutated in liver cancer or *FANCM*, known susceptibility gene for breast cancer (34) but identified here as a somatic driver.

**Figure 3.**
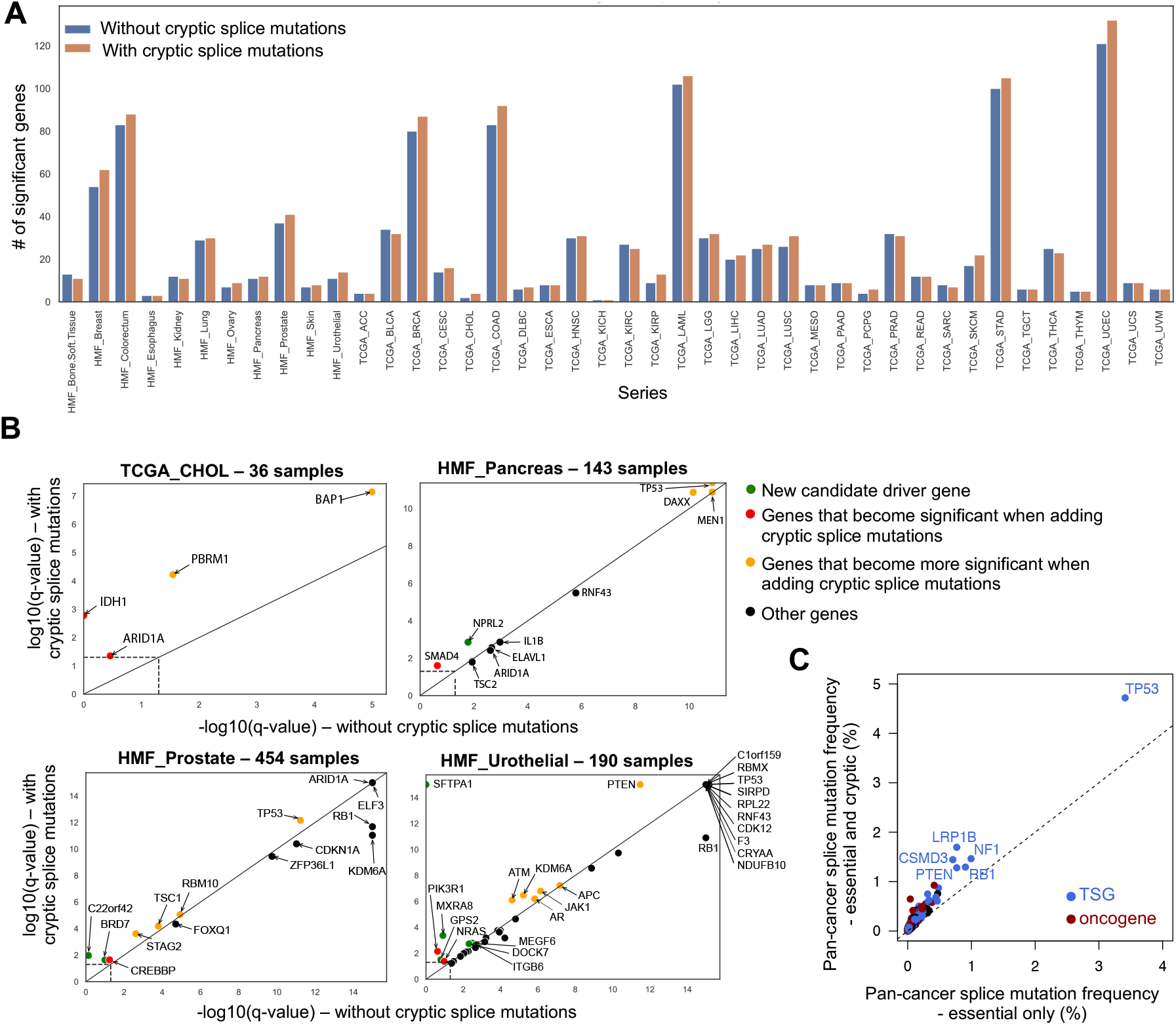
Cryptic splice mutations enhance the landscape of driver genes and mutations. (**A**) Number of significant genes identified by MutSigCV algorithm when considering only classical coding mutations (in blue) or when adding cryptic splice mutations revealed by SpliceAI (in yellow). (**B**) Comparison of MutSigCV q-values when considering only classical coding mutations or when adding cryptic splice mutations. Four series are shown as examples. Results for all series are shown in **Supplementary Figure S2**. (**C**) Frequency of splice mutations of Cancer Gene Census genes considering only essential or both essential and cryptic splice mutations. Tumor suppressor genes and oncogenes are distinguished with a color code.

In addition to finding new driver genes, cryptic splice mutations may reveal additional driver mutations in known cancer genes, incorrectly annotated as harmless variants otherwise (e.g. intronic or synonymous). To quantify this, we estimated the mutation frequency of the 695 genes from the Cancer Gene Census present in our large pan-cancer series, considering only classical coding mutations (missense, nonsense, essential splice, frameshift and in-frame indels) or adding cryptic splice mutations (**Supplementary Table S3**). On average, cryptic splice mutations increased splice mutation frequencies of driver genes by 100%, and particularly in tumor suppressor genes (**Figure 3C**). For example, TP53 mutation frequency increased from 3.4% to 4.7% when considering cryptic splice mutations. In total, we identified 7,043 cryptic splice mutations in cancer genes versus 7,475 essential splice mutations, thus doubling the prevalence of splice mutations in those genes. Considering cryptic splice mutations thus improves the power of driver gene discovery, and increases the mutation frequencies of driver genes by rescuing variants misclassified as harmless.

### Mutational processes at the origin of splice mutations

We next investigated the mutational processes at the origin of splice mutations. To that aim, we extracted the contribution of COSMIC mutational signatures (31) to each of the 2,699 tumors from the ICGC-PCAWG project. We then used Palimpsest (30) to estimate the probability of each splice mutation being due to each mutational process. By summing these probabilities over splice mutations (essential splice + cryptic splice mutations identified by SpliceAI) and other mutations, we could estimate the contribution of signatures to both mutation sets in each cancer type (**Figure 4A**). The contribution of signatures to splice and other mutations were globally consistent and highly dependent on the tumor type. For example, splice mutations were mostly attributed to single base substitution signature SBS4 (tobacco-related) in lung, wheras they were mostly attributed to SBS7a/b (UV-related) in melanoma. Yet, we identified signatures with significantly higher probabilities of giving rise to splice than other mutations (**Supplementary Table S4**). Interestingly, clock-like signatures SBS1 (due to the spontaneous deamination of mCpGs) and SBS5 (unknown etiology, age-related) were significantly enriched in splice mutations of several cancer types, including esophageal, stomach and colorectal cancers (**Figure 4B**). Several signatures associated with DNA repair defects were also enriched in splice mutations (**Figure 4C**). This was the case for the mismatch repair deficiency signatures SBS26 (uterus) and SBS44 (stomach, colorectal), as well as the homologous repair deficiency signature SBS3 (breast, ovary). This may indicate that the preferential sequence contexts of these mutational processes overlap with splicing regulatory sequences. In contrast, other signatures were significantly less prevalent in splice mutations, including SBS17b signature (associated with fluoropyrimidine-based chemotherapy) in esophageal and stomach cancer. Altogether, these data indicate that the mutational processes at the origin of splice mutations are highly cancer-dependent, and that some processes (clock-like, DNA repair deficiencies) are particularly prone to generating splice mutations.

**Figure 4.**
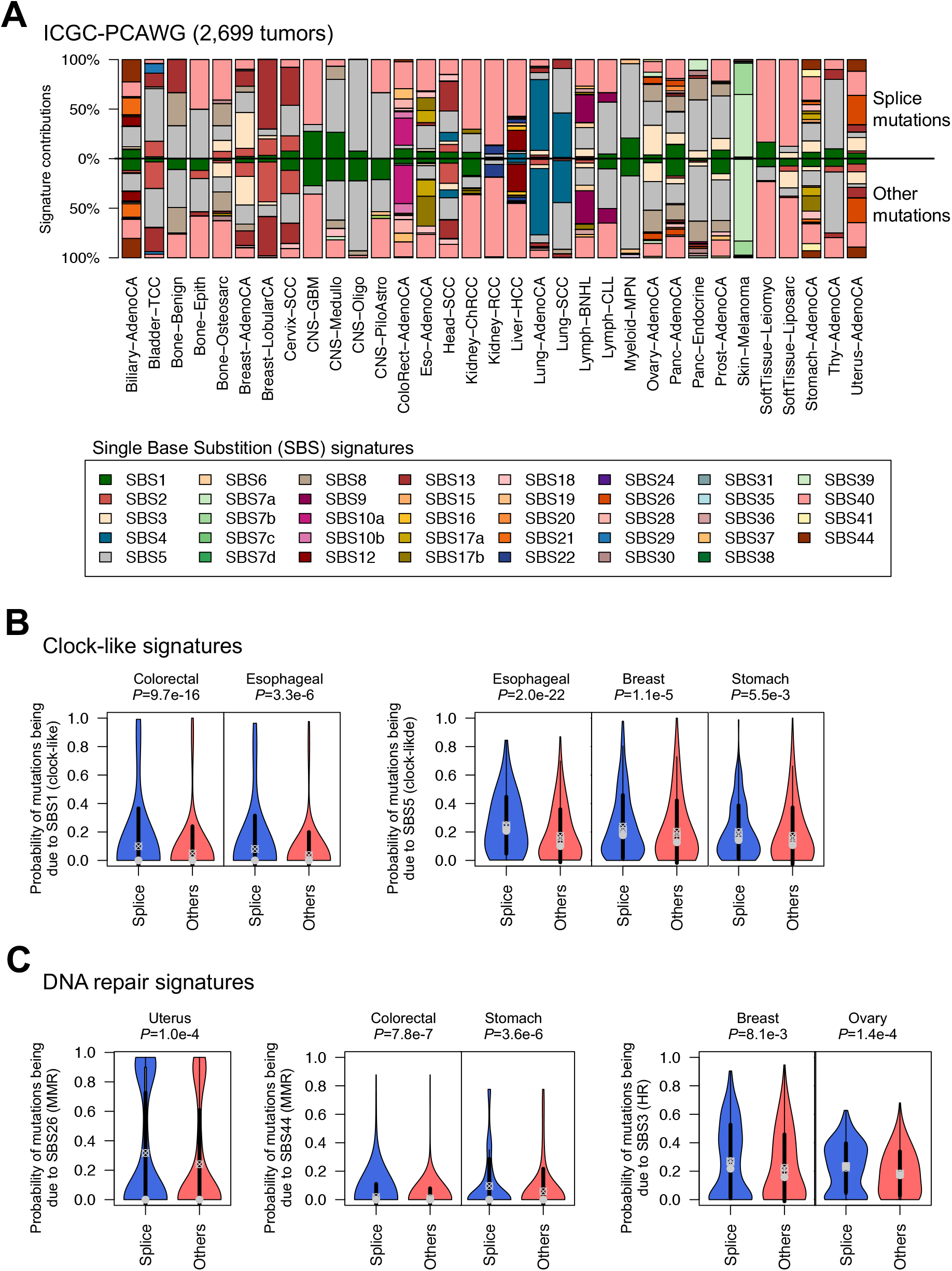
Mutational processes generating splice mutations. (**A**) Mutational signatures contributing to splice (top) and other (bottom) mutations in each series of the ICGC-PCAWG project. Splice mutations include both mutations in essential splice sites and cryptic splice mutations identified by SpliceAI (DS ≥ 0.5). (**B**) Clock-like and DNA repair signatures have significantly higher probabilities of giving rise to splice mutations compared to other mutations in several cancer types. P-values were obtained using Wilcoxon rank-sum tests.

## DISCUSSION

Our comparison of SpliceAI, GeneSplicer and MaxEntScan in a cancer setting is consistent with a previous benchmark in a context of germline variants (18), showing a higher sensitivity and specificity of SpliceAI. Of note, SpliceAI includes long sequences as input (10 kb around the mutation), which allows the tool to account for long-range sequence determinants. This may explain at least in part the higher performance of SpliceAI. In addition, SpliceAI returns predictions for mutations that can be located deeply in the introns, while GeneSplicer and MaxEntScan are restricted to mutations close to the essential splice site. Another approach, MiSplice, was recently proposed and applied to large pancancer data sets (35). While this approach unraveled and validated many cryptic splice sites, it requires as input both DNA and RNA sequencing data. By contrast, SpliceAI requires only the DNA sequence and can thus be used to annotate any variant list identified by WES/WGS.

SpliceAI identified 115,197 cryptic splice mutations in our large pan-cancer cohort, 34% are synonymous or intronic variants hence classified as low impact variants by classical variant annotation tools. We showed that rescuing these cryptic splice mutations increases the power of driver gene discovery with MutSigCV, and doubles the frequency of splice alterations in known tumor suppressor genes. Cryptic splice variants may explain part of the 5% of cancer cases without any driver alteration identified (26). In addition, 28% of cryptic splice mutations are annotated as missense variants. These mutations may have dual consequences, leading to missense and/or splice alteration depending on the RNA molecule, exemplified by the ARID1A mutation displayed in **Figure 1G**.

Splice mutations may occur in different sequence contexts and can be generated by distinct mutational processes depending on the cancer type. Yet, we identified signatures enriched in splice mutations, related to clock-like processes, mismatch repair and homologous repair deficiencies. These associations may result from an overrepresentation of the sequence contexts in which these processes are particularly active in splice regulatory elements. The contribution of DNA repair signatures to splice mutations may lead to a higher risk of acquiring splice mutations in patients carrying defects in DNA repair enzymes.

This study highlights the power of deep learning to uncover previously overlooked regulatory mutations in cancer sequencing data, and emphasizes the importance of better annotating cryptic splice mutations in research series and clinical sequencing data.

## AVAILABILITY

LICA-FR data used in this manuscript are hosted in the European Genome-Phenome Archive (https://ega-archive.org). We downloaded public cancer genomic data from the ICGC-PCAWG (https://dcc.icgc.org/pcawg), TCGA (https://gdc.cancer.gov/about-data/publications/pancanatlas) and HMF (https://www.hartwigmedicalfoundation.nl/en/data/database/) websites. The pyRNA library that we developed to mine RNA-seq data in Python is available on GitHub (https://github.com/FunGeST/pyRNA).

## SUPPLEMENTARY DATA

The following supplementary data are provided:

**Supplementary Figures** (pdf file)

**Supplementary Figure S1:** Proportion of splicing alteration types and VEP annotations according to SpliceAI DS scores

**Supplementary Figure S2:** MutSigCV q-values including or not cryptic splice mutations for each HMF and TCGA series

**Supplementary Tables** (Excel file)

**Supplementary Table S1:** Cryptic splice mutations identified by SpliceAI, MaxEntScan and GeneSplicer in the LICA-FR series.

**Supplementary Table S2:** 153 extra-significant genes identified by MutSigCV when including cryptic splice mutations.

**Supplementary Table S3:** Mutation rates of Cancer Gene Census genes in a large pan-cancer dataset

**Supplementary Table S4:** Mutational signatures contributing preferentially to splice mutations.

## ACKNOWLEDGEMENT

We thank members of the thesis committee of Raphaël Teboul (Tatiana Popova, Andrei Zinovyev, Josh Waterfall and Stéphane Minvielle) for fruitful discussions and advice.

## REFERENCES

1. Black, D.L. (2003) Mechanisms of alternative pre-messenger RNA splicing. Annu Rev Biochem, 72, 291–336.

2. Pan, Q., Shai, O., Lee, L.J., Frey, B.J. and Blencowe, B.J. (2008) Deep surveying of alternative splicing complexity in the human transcriptome by high-throughput sequencing. Nat Genet, 40, 1413–1415.

3. Romero, P.R., Zaidi, S., Fang, Y.Y., Uversky, V.N., Radivojac, P., Oldfield, C.J., Cortese, M.S., Sickmeier, M., LeGall, T., Obradovic, Z., et al. (2006) Alternative splicing in concert with protein intrinsic disorder enables increased functional diversity in multicellular organisms. Proc Natl Acad Sci U S A, 103, 8390–8395.

4. Wang, Z. and Burge, C.B. (2008) Splicing regulation: From a parts list of regulatory elements to an integrated splicing code. RNA, 14, 802–813.

5. Sterne-Weiler, T. and Sanford, J.R. (2014) Exon identity crisis: disease-causing mutations that disrupt the splicing code. Genome Biology, 15, 201.

6. Xiong, H.Y., Alipanahi, B., Lee, L.J., Bretschneider, H., Merico, D., Yuen, R.K.C., Hua, Y., Gueroussov, S., Najafabadi, H.S., Hughes, T.R., et al. (2015) The human splicing code reveals new insights into the genetic determinants of disease. Science, 347, 1254806.

7. Scotti, M.M. and Swanson, M.S. (2016) RNA mis-splicing in disease. Nat Rev Genet, 17, 19–32.

8. Anczuków, O. and Krainer, A.R. (2016) Splicing-factor alterations in cancers. RNA, 22, 1285–1301.

9. Sveen, A., Kilpinen, S., Ruusulehto, A., Lothe, R.A. and Skotheim, R.I. (2016) Aberrant RNA splicing in cancer; expression changes and driver mutations of splicing factor genes. Oncogene, 35, 2413–2427.

10. Sondka, Z., Bamford, S., Cole, C.G., Ward, S.A., Dunham, I. and Forbes, S.A. (2018) The COSMIC Cancer Gene Census: describing genetic dysfunction across all human cancers. Nat Rev Cancer, 18, 696–705.

11. Burge, C. and Karlin, S. (1997) Prediction of complete gene structures in human genomic DNA. J Mol Biol, 268, 78–94.

12. Pertea, M., Lin, X. and Salzberg, S.L. (2001) GeneSplicer: a new computational method for splice site prediction. Nucleic Acids Res, 29, 1185–1190.

13. Cai, D., Delcher, A., Kao, B. and Kasif, S. (2000) Modeling splice sites with Bayes networks. Bioinformatics, 16, 152–158.

14. Arita, M., Tsuda, K. and Asai, K. (2002) Modeling splicing sites with pairwise correlations. Bioinformatics, 18 Suppl 2, S27–34.

15. Yeo, G. and Burge, C.B. (2004) Maximum entropy modeling of short sequence motifs with applications to RNA splicing signals. J Comput Biol, 11, 377–394.

16. Shamsani, J., Kazakoff, S.H., Armean, I.M., McLaren, W., Parsons, M.T., Thompson, B.A., O’Mara, T.A., Hunt, S.E., Waddell, N. and Spurdle, A.B. (2019) A plugin for the Ensembl Variant Effect Predictor that uses MaxEntScan to predict variant spliceogenicity. Bioinformatics, 35, 2315–2317.

17. Reese, M.G., Eeckman, F.H., Kulp, D. and Haussler, D. (1997) Improved splice site detection in Genie. J Comput Biol, 4, 311–323.

18. Jaganathan, K., Kyriazopoulou Panagiotopoulou, S., McRae, J.F., Darbandi, S.F., Knowles, D., Li, Y.I., Kosmicki, J.A., Arbelaez, J., Cui, W., Schwartz, G.B., et al. (2019) Predicting Splicing from Primary Sequence with Deep Learning. Cell, 176, 535–548.e24.

19. Schulze, K., Imbeaud, S., Letouzé, E., Alexandrov, L.B., Calderaro, J., Rebouissou, S., Couchy, G., Meiller, C., Shinde, J., Soysouvanh, F., et al. (2015) Exome sequencing of hepatocellular carcinomas identifies new mutational signatures and potential therapeutic targets. Nat. Genet., 47, 505–511.

20. Letouzé, E., Shinde, J., Renault, V., Couchy, G., Blanc, J.-F., Tubacher, E., Bayard, Q., Bacq, D., Meyer, V., Semhoun, J., et al. (2017) Mutational signatures reveal the dynamic interplay of risk factors and cellular processes during liver tumorigenesis. Nat Commun, 8, 1315.

21. Bayard, Q., Meunier, L., Peneau, C., Renault, V., Shinde, J., Nault, J.-C., Mami, I., Couchy, G., Amaddeo, G., Tubacher, E., et al. (2018) Cyclin A2/E1 activation defines a hepatocellular carcinoma subclass with a rearrangement signature of replication stress. Nat Commun, 9, 5235.

22. Hirsch, T.Z., Negulescu, A., Gupta, B., Caruso, S., Noblet, B., Couchy, G., Bayard, Q., Meunier, L., Morcrette, G., Scoazec, J.-Y., et al. (2020) BAP1 mutations define a homogeneous subgroup of hepatocellular carcinoma with fibrolamellar-like features and activated PKA. J. Hepatol., 72, 924–936.

23. Péneau, C., Imbeaud, S., La Bella, T., Hirsch, T.Z., Caruso, S., Calderaro, J., Paradis, V., Blanc, J.-F., Letouzé, E., Nault, J.-C., et al. (2021) Hepatitis B virus integrations promote local and distant oncogenic driver alterations in hepatocellular carcinoma. Gut, 10.1136/gutjnl-2020-323153.

24. Hirsch, T.Z., Pilet, J., Morcrette, G., Roehrig, A., Monteiro, B.J.E., Molina, L., Bayard, Q., Trépo, E., Meunier, L., Caruso, S., et al. (2021) Integrated Genomic Analysis Identifies Driver Genes and Cisplatin-Resistant Progenitor Phenotype in Pediatric Liver Cancer. Cancer Discov, 11, 2524–2543.

25. Molina, L., Zhu, J., Trépo, E., Bayard, Q., Amaddeo, G., GENTHEP Consortium, Blanc, J.-F., Calderaro, J., Ma, X., Zucman-Rossi, J., et al. (2022) Bi-allelic hydroxymethylbilane synthase inactivation defines a homogenous clinico-molecular subtype of hepatocellular carcinoma. J Hepatol, 77, 1038–1046.

26. Campbell, P.J., Getz, G., Korbel, J.O., Stuart, J.M., Jennings, J.L., Stein, L.D., Perry, M.D., Nahal-Bose, H.K., Ouellette, B.F.F., Li, C.H., et al. (2020) Pan-cancer analysis of whole genomes. Nature, 578, 82–93.

27. Priestley, P., Baber, J., Lolkema, M.P., Steeghs, N., de Bruijn, E., Shale, C., Duyvesteyn, K., Haidari, S., van Hoeck, A., Onstenk, W., et al. (2019) Pan-cancer whole-genome analyses of metastatic solid tumours. Nature, 575, 210–216.

28. Hoadley, K.A., Yau, C., Hinoue, T., Wolf, D.M., Lazar, A.J., Drill, E., Shen, R., Taylor, A.M., Cherniack, A.D., Thorsson, V., et al. (2018) Cell-of-Origin Patterns Dominate the Molecular Classification of 10,000 Tumors from 33 Types of Cancer. Cell, 173, 291–304.e6.

29. Lawrence, M.S., Stojanov, P., Polak, P., Kryukov, G.V., Cibulskis, K., Sivachenko, A., Carter, S.L., Stewart, C., Mermel, C.H., Roberts, S.A., et al. (2013) Mutational heterogeneity in cancer and the search for new cancer-associated genes. Nature, 499, 214–218.

30. Shinde, J., Bayard, Q., Imbeaud, S., Hirsch, T.Z., Liu, F., Renault, V., Zucman-Rossi, J. and Letouzé, E. (2018) Palimpsest: an R package for studying mutational and structural variant signatures along clonal evolution in cancer. Bioinformatics, 34, 3380–3381.

31. Alexandrov, L.B., Kim, J., Haradhvala, N.J., Huang, M.N., Tian Ng, A.W., Wu, Y., Boot, A., Covington, K.R., Gordenin, D.A., Bergstrom, E.N., et al. (2020) The repertoire of mutational signatures in human cancer. Nature, 578, 94–101.

32. Spain, B.H., Bowdish, K.S., Pacal, A.R., Staub, S.F., Koo, D., Chang, C.Y., Xie, W. and Colicelli, J. (1996) Two human cDNAs, including a homolog of Arabidopsis FUS6 (COP11), suppress G-protein- and mitogen-activated protein kinase-mediated signal transduction in yeast and mammalian cells. Mol Cell Biol, 16, 6698–6706.

33. Jamialahmadi, O., Mancina, R.M., Ciociola, E., Tavaglione, F., Luukkonen, P.K., Baselli, G., Malvestiti, F., Thuillier, D., Raverdy, V., Männistö, V., et al. (2021) Exome-Wide Association Study on Alanine Aminotransferase Identifies Sequence Variants in the GPAM and APOE Associated With Fatty Liver Disease. Gastroenterology, 160, 1634–1646.e7.

34. Kiiski, J.I., Pelttari, L.M., Khan, S., Freysteinsdottir, E.S., Reynisdottir, I., Hart, S.N., Shimelis, H., Vilske, S., Kallioniemi, A., Schleutker, J., et al. (2014) Exome sequencing identifies FANCM as a susceptibility gene for triple-negative breast cancer. Proceedings of the National Academy of Sciences, 111, 15172–15177.

35. Cao, S., Zhou, D.C., Oh, C., Jayasinghe, R.G., Zhao, Y., Yoon, C.J., Wyczalkowski, M.A., Bailey, M.H., Tsou, T., Gao, Q., et al. (2020) Discovery of driver non-coding splice-site-creating mutations in cancer. Nat Commun, 11, 5573.

