## Supplementary Figure for "Discovering cryptic splice mutations in cancers via a deep neural network framework"

### Supplementary Data

#### List of Supplementary Figures

**Supplementary Figure S1:** Proportion of splicing alteration types and VEP annotations according to SpliceAI DS scores

**Supplementary Figure S2:** MutSigCV q-values including or not cryptic splice mutations for each HMF and TCGA series

#### List of Supplementary Tables (provided as Excel files)

**Supplementary Table S1:** Cryptic splice mutations identified by SpliceAI, MaxEntScan and GeneSplicer in the LICA-FR series.

**Supplementary Table S2:** 153 extra-significant genes identified by MutSigCV when including cryptic splice mutations.

**Supplementary Table S3:** Mutation rates of Cancer Gene Census genes in a large pan-cancer dataset

**Supplementary Table S4:** Mutational signatures contributing preferentially to splice mutations.

A

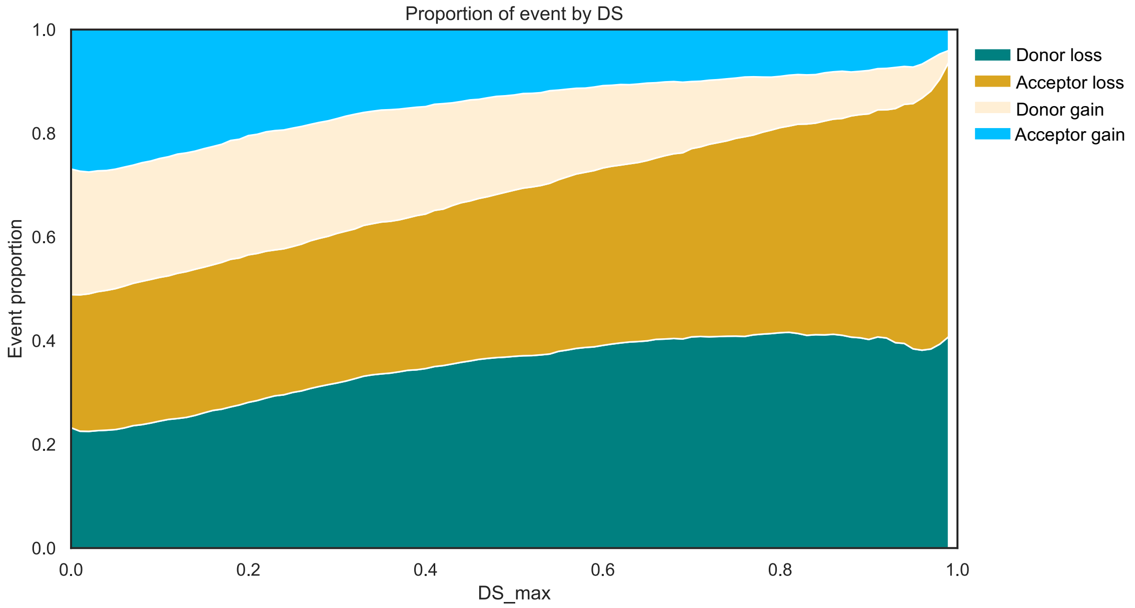


B

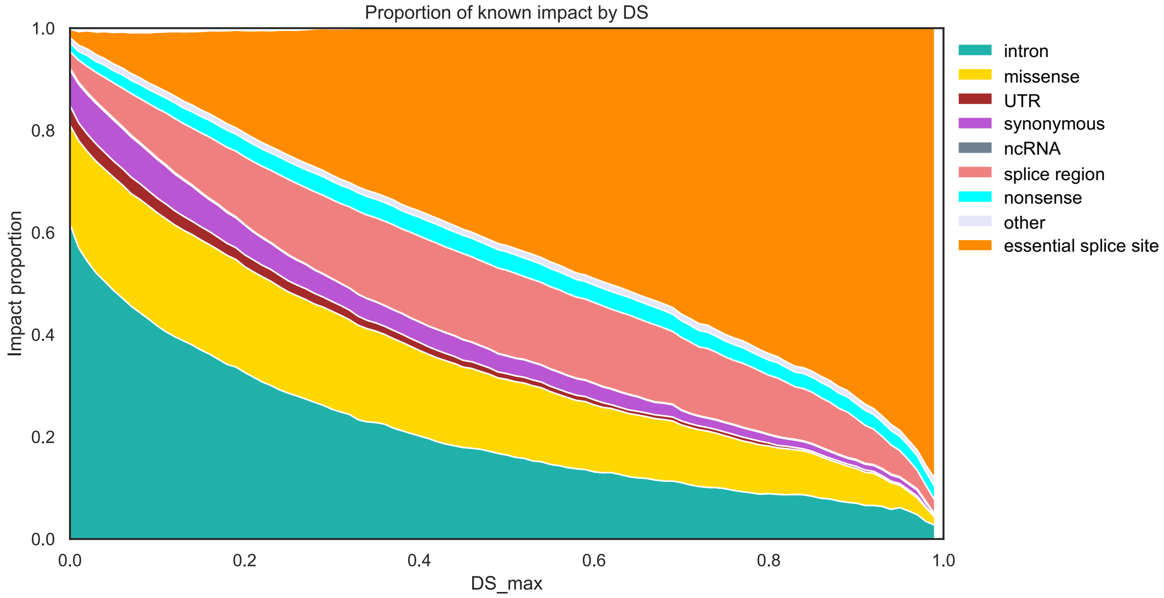


**Supplementary Figure S1: Proportion of splicing alteration types and VEP annotations according to SpliceAI DS scores. A** Proportion of donor loss (DL), acceptor loss (AL), donor gain (DG) and acceptor gain (AG) events as a function of SpliceAI delta score (DS). **B** Proportion of mutation categories (annotated by the Variant Effect Predictor) as a function of SpliceAI delta score (DS).

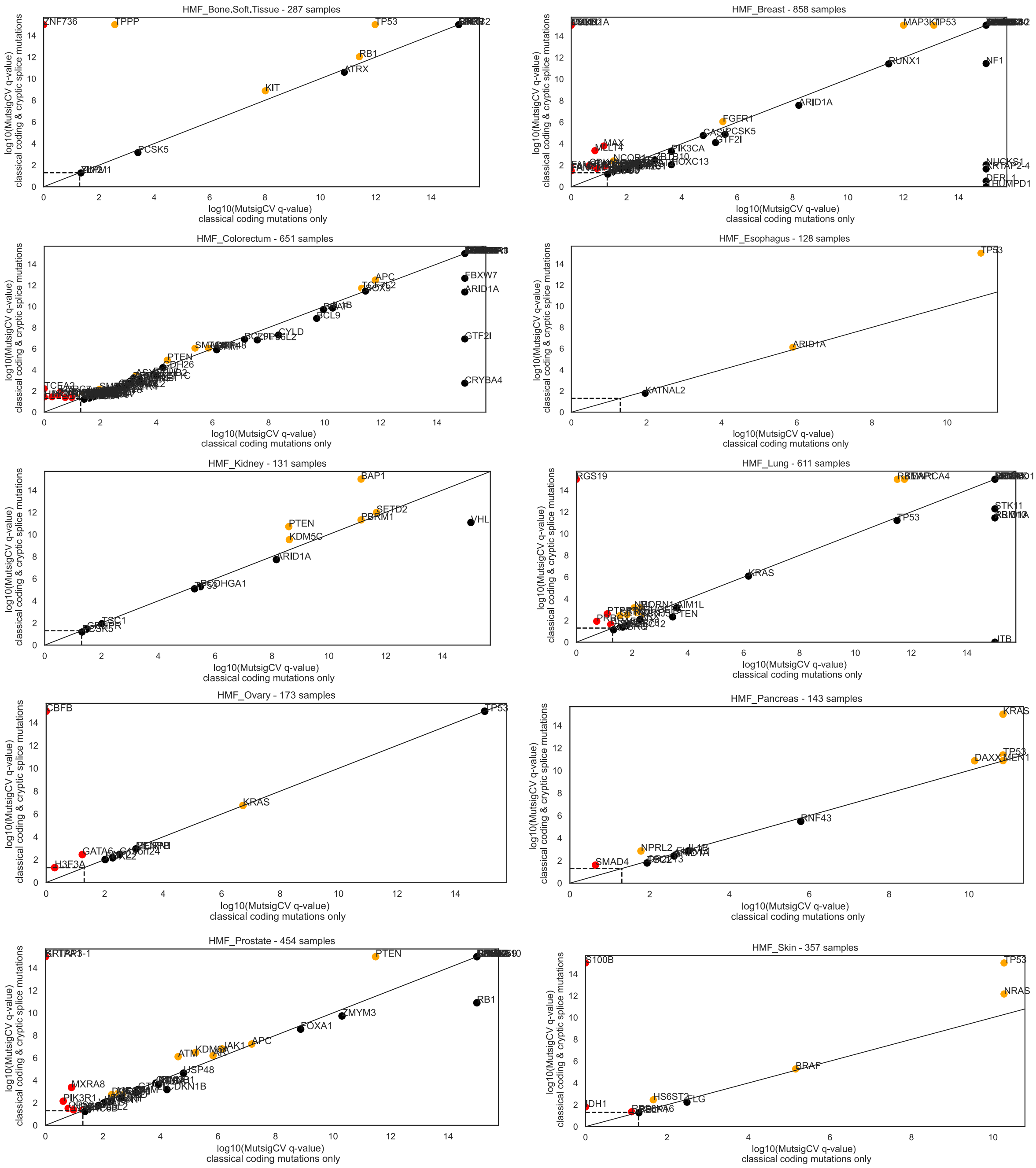

HMF_Bone.Soft.Tissue - 287 samples
ZNF736
TPPP
TP53
RB1
ATRX
KIT
PCSK5
HMF_Breast - 858 samples
MAP3K1
TP53
RUNX1
NF1
ARID1A
FGFR1
PCSK5
GTF2I
PIK3CA
MAX
HOXC13
NUCKS1
KRTAP2-4
HMF_Colorectum - 651 samples
APC
FBXW7
ARID1A
GTF2I
CRYBA4
CYLD
PTEN
BCL9
TCEA2
HMF_Esophagus - 128 samples
TP53
ARID1A
KATNAL2
HMF_Kidney - 131 samples
BAP1
SETD2
PBRM1
VHL
PTEN
KDM5C
ARID1A
TSC1
HMF_Lung - 611 samples
RGS19
SMARCA4
STK11
TP53
KRAS
PTEN
HMF_Ovary - 173 samples
CBFB
TP53
KRAS
GATA6
H3F3A
HMF_Pancreas - 143 samples
KRAS
TP53
DAXX
MEN1
RNF43
NPRL2
SMAD4
HMF_Prostate - 454 samples
PTEN
RB1
ZMYM3
FOXA1
APC
JAK1
ATM
USP48
MXRA8
PIK3R1
HMF_Skin - 357 samples
S100B
TP53
NRAS
BRAF
HS6ST2
FLG
IDH1
log10(MutsigCV q-value)
classical coding mutations only
log10(MutsigCV q-value)
classical coding & cryptic splice mutations

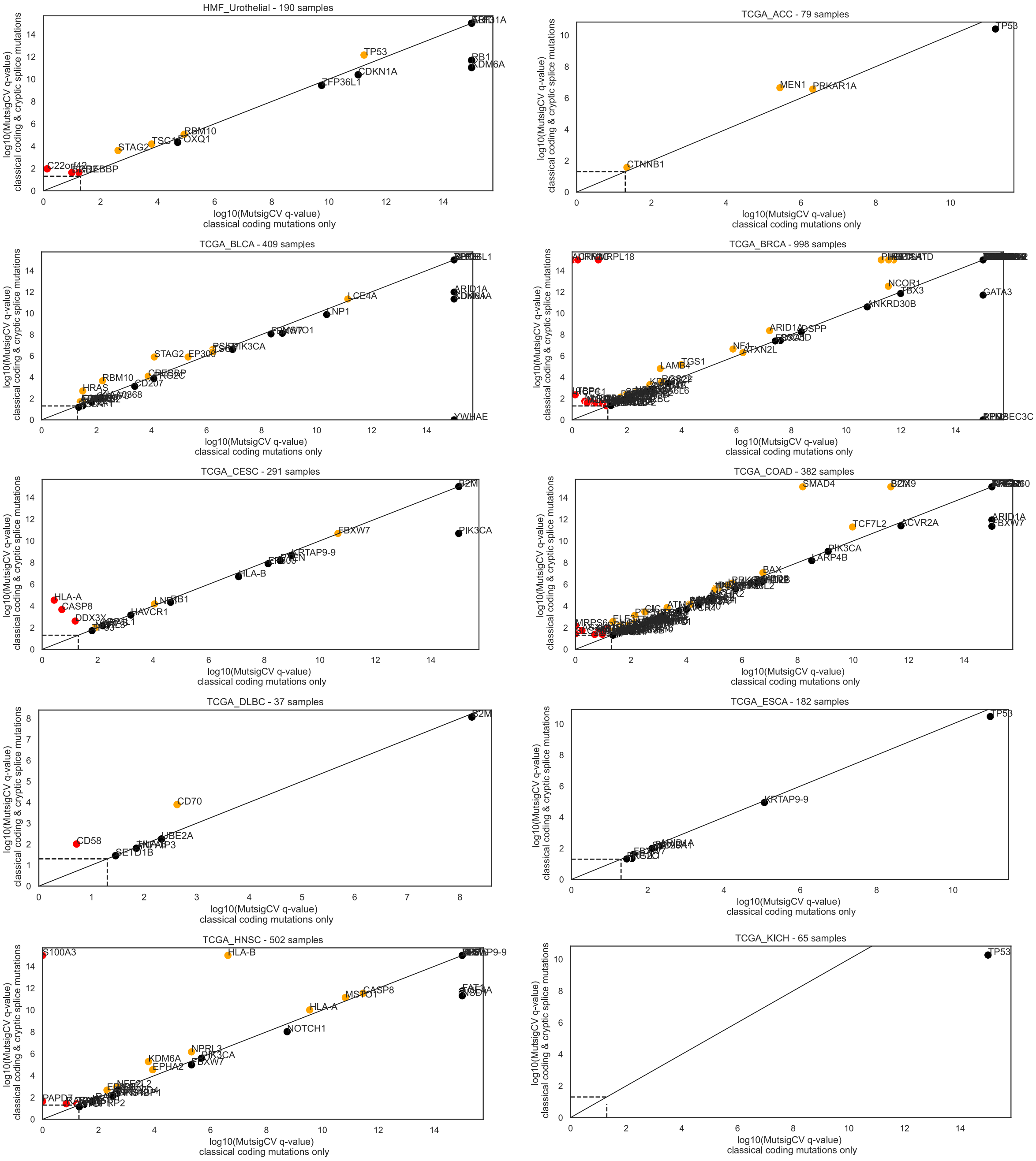

HMF_Urothelial - 190 samples
log10(MutsigCV q-value) classical coding & cryptic splice mutations
log10(MutsigCV q-value) classical coding mutations only
TP53
RB1
KDM6A
CDKN1A
ZFP36L1
RBM10
FOXQ1
TSC1
STAG2
C22orf42
TCGA_ACC - 79 samples
TP53
MEN1
PRKAR1A
CTNNB1
TCGA_BLCA - 409 samples
ARID1A
LCE4A
ELF3
PIK3CA
STAG2
EP300
RBM10
HRAS
CD207
YWHAE
TCGA_BRCA - 998 samples
MRPL18
NCOR1
FBX3
GATA3
ANKRD30B
ARID1A
DSPP
NF1
ATXN2L
LAMB4
TGS1
APOBEC3C
TCGA_CESC - 291 samples
B2M
PIK3CA
FBXW7
KRTAP9-9
HLA-B
HLA-A
CASP8
DDX3X
HAVCR1
RB1
TCGA_COAD - 382 samples
SMAD4
SOX9
TCF7L2
ACVR2A
ARID1A
FBXW7
PIK3CA
LARP4B
BAX
ATM
TCGA_DLBC - 37 samples
B2M
CD70
CD58
UBE2A
SETD1B
TCGA_ESCA - 182 samples
TP53
KRTAP9-9
TCGA_HNSC - 502 samples
S100A3
HLA-B
KRTAP9-9
FAT1
CASP8
MSTO1
HLA-A
NOTCH1
NPRL3
PIK3CA
KDM6A
FBXW7
EPHA2
NFE2L2
PAPD7
TCGA_KICH - 65 samples
TP53

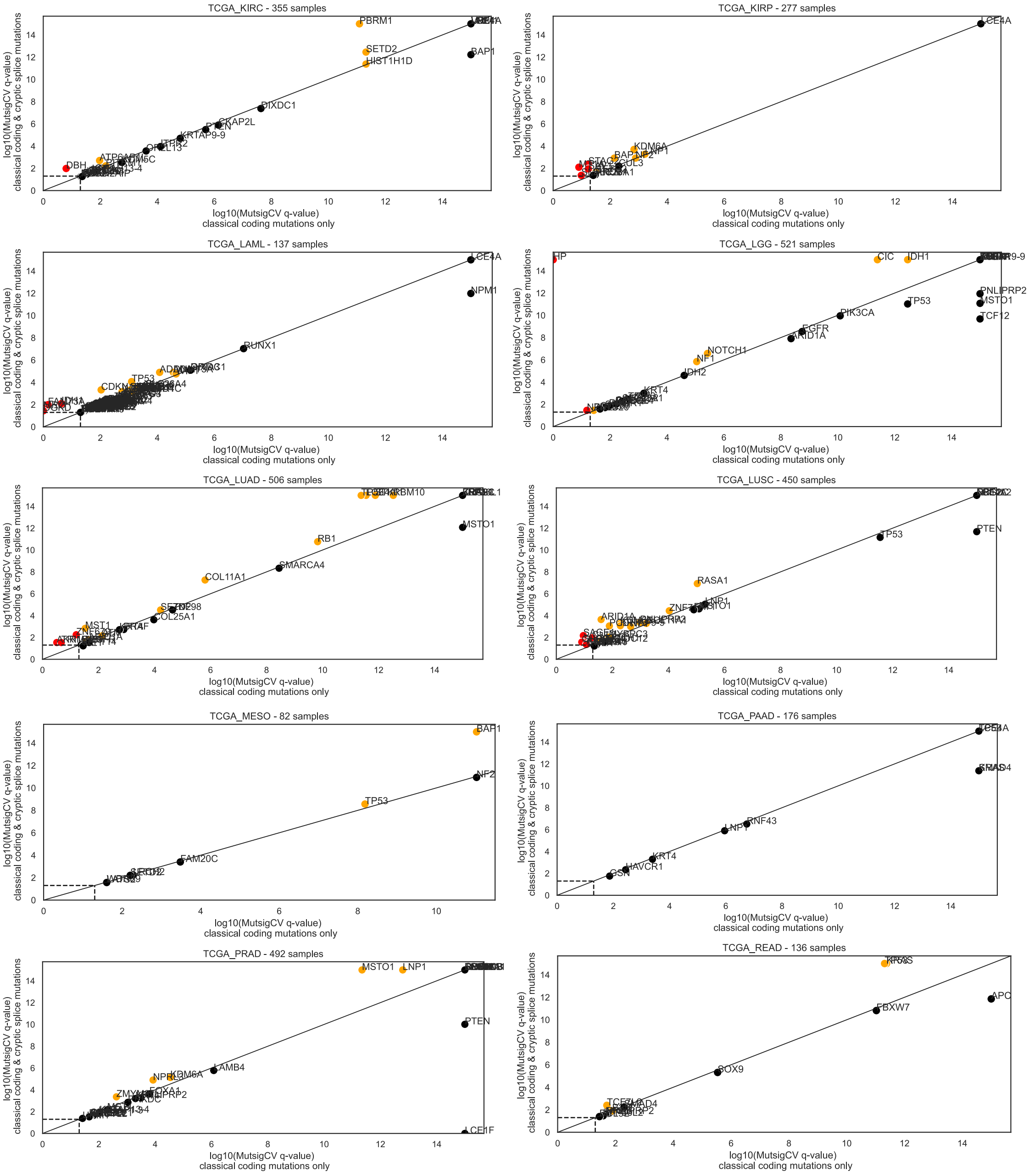

TCGA_KIRC - 355 samples
PBRM1
SETD2
HIST1H1D
BAP1
DIXDC1
CKAP2L
KRTAP9-9
DBH
TCGA_KIRP - 277 samples
LCE4A
KDM6A
CUL3
TCGA_LAML - 137 samples
LCE4A
NPM1
RUNX1
TP53
TCGA_LGG - 521 samples
HP
CIC
IDH1
PNLIPRP2
MSTO1
TCF12
TP53
PIK3CA
EGFR
ARID1A
NOTCH1
NF1
IDH2
KRT4
TCGA_LUAD - 506 samples
RBM10
MSTO1
RB1
SMARCA4
COL11A1
COL25A1
MST1
TCGA_LUSC - 450 samples
TP53
PTEN
RASA1
LNP1
MSTO1
ARID1A
TCGA_MESO - 82 samples
BAP1
NF2
TP53
FAM20C
TCGA_PAAD - 176 samples
RNF43
KRT4
HAVCR1
TCGA_PRAD - 492 samples
MSTO1
LNP1
PTEN
LAMB4
KDM6A
LCE1F
TCGA_READ - 136 samples
APC
FBXW7
SOX9
log10(MutsigCV q-value)
classical coding mutations only
log10(MutsigCV q-value)
classical coding & cryptic splice mutations

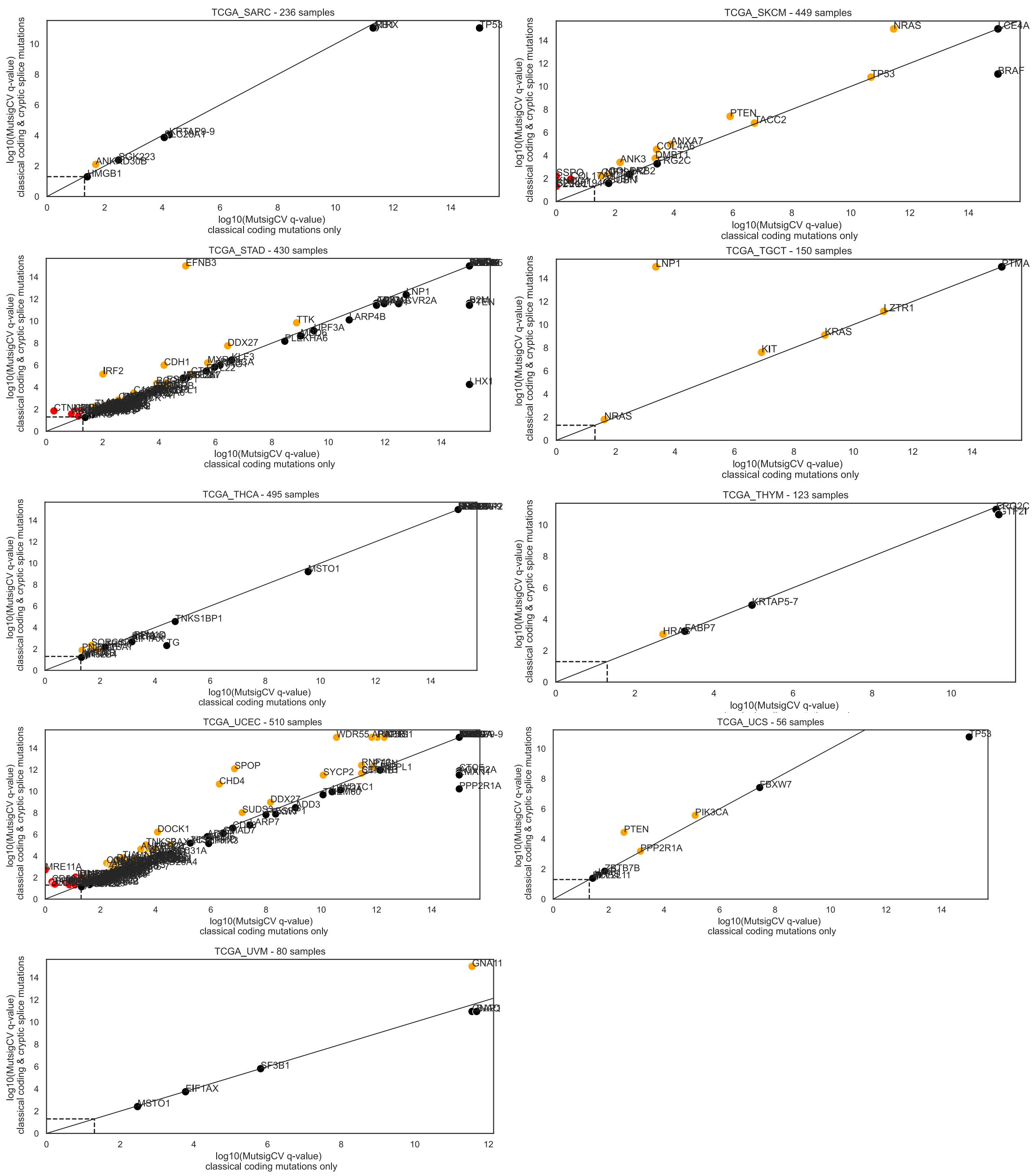


**Supplementary Figure S2: MutSigCV q-values including or not cryptic splice mutations for each HMF and TCGA series.** MutSigCV q-values were calculated for each HMF/TCGA series considering only classical coding and essential splice mutations, or when adding cryptic splice mutations. Genes significant in one or the other test are represented with a color code indicating whether they become significant or increase their significance level when adding cryptic splice mutations.
